# Impact of Genome Architecture Upon the Functional Activation and Repression of *Hox* Regulatory Landscapes

**DOI:** 10.1101/587303

**Authors:** Eddie Rodríguez-Carballo, Lucille Lopez-Delisle, Nayuta Yakushiji-Kaminatsui, Asier Ullate-Agote, Denis Duboule

## Abstract

The spatial organization of the mammalian genome relies upon the formation of chromatin domains of various scales. At the level of gene regulation *in cis*, collections of enhancer sequences define large regulatory landscapes that usually match with the presence of topologically associating domains (TADs). These domains are largely determined by bound CTCF molecules and often contain ranges of enhancers displaying similar or related tissue specificity, suggesting that in some cases such domains may act as coherent regulatory units, with a global on or off state.

**Results:** By using the *HoxD* gene cluster as a paradigm, we investigated the effect of large genomic rearrangements affecting the two TADs flanking this locus, including their fusion into a single chromatin domain. We show that, within a single hybrid TAD, the activation of both proximal and distal limb enhancers initially positioned in either TADs globally occurred as when both TADs are intact. We also show that the timely implementation of distal limb enhancers depends on whether or not target genes had previously responded to proximal enhancers, due to the presence or absence of H3K27me3 marks.

**Conclusions:** From this work, we conclude that antagonistic limb proximal and distal enhancers can exert their specificities when positioned into the same TAD and in the absence of their genuine target genes. We also conclude that removing these target genes reduced the coverage of a regulatory landscape by chromatin marks associated with silencing and thus prolonged its activity in time. Since Polycomb group proteins are mainly recruited at the *Hox* gene cluster, our results suggest that Polycomb Repressive Complex 2 (PRC2) can extend its coverage to far-*cis* regulatory sequences as long as confined to the neighboring TAD structure.

## BACKGROUND

Attempts to understand the spatial organization of the genome in the nucleus have recently led to models accounting for the relationship between genome structure and gene regulation (see [1]). The development of chromosome conformation capture techniques associated with deep sequencing has thus allowed the resolution of DNA interactions at a small scale [2]. These interactions can be either structural or functional, i.e. they can be present regardless of the transcriptional outcome or alternatively, they can fluctuate according to cell-type specific context depending upon the transcriptional status [3]. Constitutive contacts generally tend to fit the loop extrusion model, whereby the packed network of chromatin loops would form as a result of DNA extrusion by an ATP-dependent cohesin-based complex. The loops are stabilized whenever this cohesin ring meets two CTCF molecules bound with convergent orientations [4–6].

Chromatin is organized in several levels of interactions, loops and domains. At the level of gene regulation, topologically associating domains (TADs) [7, 8][9] usually match large domains of long-range gene regulation referred to as regulatory landscapes [10]. These structures are globally detected in all cell types and conserved across vertebrate species [7, 11–15]. The experimental depletion of either CTCF or cohesin subunits lead to a loss of both loop organization and TAD structure. Under these conditions, however, the effects upon gene transcription were limited and the formation of larger structures (compartments), which may also be functionally relevant, still occurred although in an altered manner [16–19][20].

Compartments contain chromatin domains labelled by various epigenetic marks. Inactive chromatin domains labelled by histone H3 lysine 27 trimethylation (H3K27me3), resulting from the presence of Polycomb group protein complexes, have been associated either with compartment A [21] or with a compartment B1, distinct from the genuine heterochromatin B compartment [5], which may segregate from other chromatin domains through phase separation [22, 23]. In addition, facultative heterochromatin (H3K27me3-positive) was shown to correlate with long-distance interactions either in stem cells [24–26] or during embryonic development [21, 27].

Distinct functional states associated with various chromatin structures are not as clear when TADs are considered. While several examples exist showing the functional coherence of multiple enhancer sequences present within one particular TAD [28–31][32], the definition of TADs as global independent regulatory units still lacks experimental evidence. In many instances indeed, TADs include either series of enhancers with the same -or related-specificity, or enhancers with distinct tissue-specific potentials but involved in the pleiotropic regulation of the same target gene(s). However, whether or not the entire TAD adopts a global on or off state, for example related to a particular architecture, remains to be established.

A useful experimental paradigm to address this question is the mammalian *HoxD* gene cluster, a group of genes located at the intersection between two TADs displaying distinct functional specificities [33]. During limb development, enhancers in the telomeric TAD (T-DOM) regulate the transcription of *Hoxd8* to *Hoxd11* in proximal limb bud cells. Subsequently, enhancers in the centromeric TAD (C-DOM) control from *Hoxd9* to *Hoxd13* in distal limb bud cells [33]. These different sets of target genes responding to either one of the regulatory domains are determined by a robust boundary, centered around *Hoxd11* and relying upon a collection of bound CTCF sites. Genetic analyses *in vivo* revealed that this boundary was very resilient and that even a full deletion of the gene cluster was unable to merge both TADs into one single domain, likely due to a few remaining occupied CTCF sites [34].

The analysis of different developmental contexts where *Hoxd* genes are transcribed demonstrates that these two TADs are functionally exclusive from one another, i.e. the concomitant function of enhancers belonging to the two domains has not been observed thus far. This is due to the fact that the main gene responding to C-DOM enhancers is *Hoxd13*, whose product, along with that of *Hoxa13*, has a negative effect over T-DOM enhancers through direct binding, as observed in ChIP-seq experiments [32, 35]. This bimodal regulation can also be followed by the appearance of relevant chromatin marks: while T-DOM is largely covered by H3K27ac marks in proximal limb bud cells, it becomes rapidly decorated by H3K27me3 marks at the time C-DOM starts to be active in distal cells and to accumulate H3K27ac labelling [33]. Therefore, in distal cells, not only the *Hoxd1* to *Hoxd8* gene are covered by H3K27me3 (they are no longer transcribed), but also large DNA intervals within T-DOM, reflecting the off state of this regulatory landscape and re-enforcing the idea that it may behave as a coherent regulatory unit.

In this paper, we challenged this hypothesis by investigating the effects of combining the two TADs into a single domain (a neoTAD), after deletion of a large piece of DNA containing the *HoxD* cluster as well as other boundary elements. After fusion, this neoTAD regroups enhancers that do not normally function in the same cellular context. We asked whether these various enhancers would keep their initial functional specificities or, alternatively, if they would all be active or repressed concomitantly as a result of this new topological proximity. We also used a set of inversions, which disconnected the target genes from their TADs to evaluate the functional and epigenetic behavior of regulatory sequences in the absence of their target genes.

## RESULTS

In order to better visualize the spatial distribution of the two TADs associated with the *HoxD* cluster (Fig. 1A), we modeled their structures in 3D by using Hi-C matrices [34] for both distal and proximal E12.5 limb bud cells (Fig. 1B) and the TADkit scripts package as a 3D modeling viewer [36]. In the wild-type condition, the *HoxD* cluster contained a strong boundary and was thus positioned between the two regulatory domains T-DOM and C-DOM, in both distal and proximal limb cells (Fig. 1B). In both tissues, the region called CS38-41 (Fig.1, red disk) established a weaker boundary between two sub-TADs in T-DOM. The structure and separation between the two regulatory domains were generally conserved between the two cell types, although with some minor differences.

**Fig. 1.**
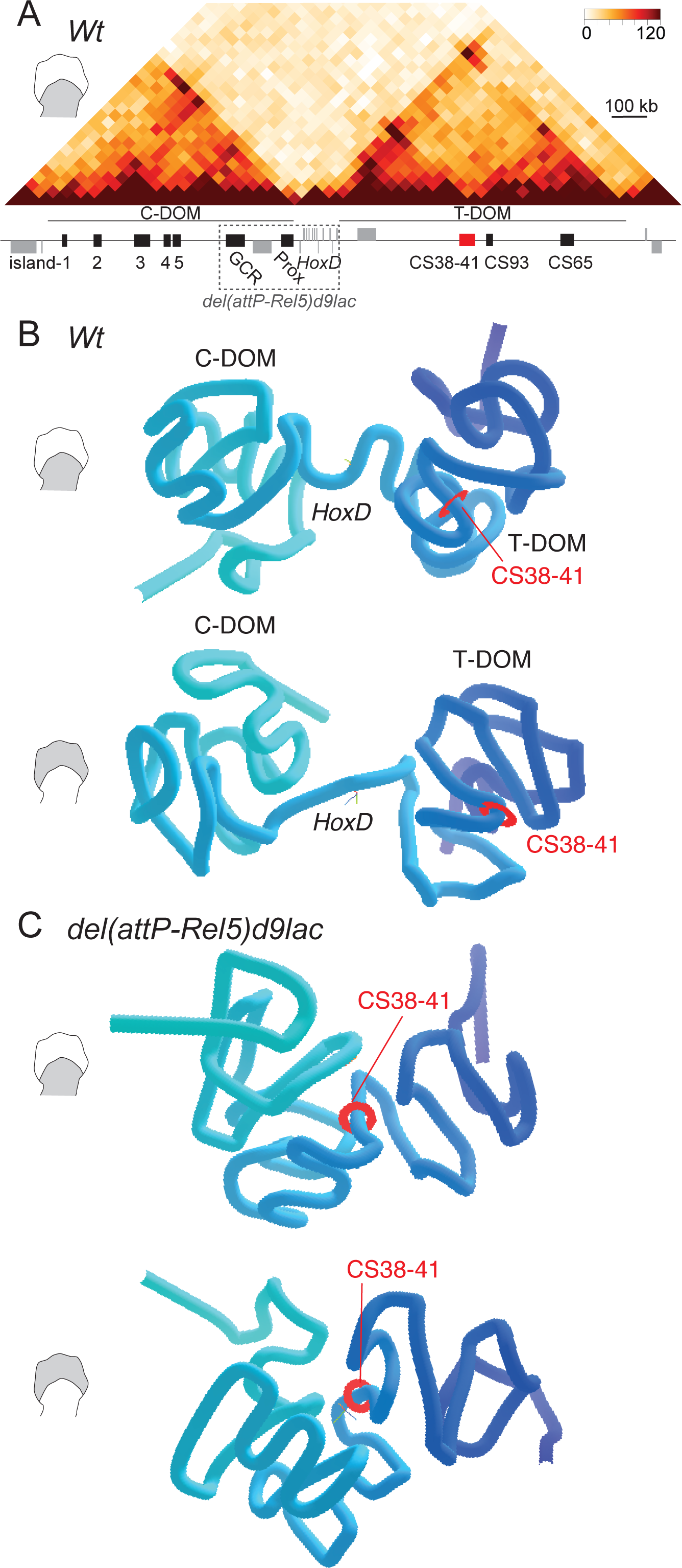
3D-representation of the *HoxD* locus in control (*Wt*) and mutant limb buds. (*A*) Hi-C map showing the distribution of TADs on either side of the *HoxD* locus in proximal limb and its associated genes (grey boxes) and regulatory regions (black and red boxes). The dashed rectangle illustrates the deletion in the *del(attP-Rel5)d9lac* allele. (*B*) Three-dimensional modeling of *HoxD* associated TADs derived from Hi-C datasets obtained from wild type (*Wt*) proximal (top) and distal (bottom) limb bud cells (schemes on the left). (*C*) Comparative modeling from the *del(attP-Rel5)d9lac* mutant proximal (top) and distal (bottom) limb bud cells. The red disk shows the position of region CS38-41 to be used as a reference point in all representations. T-DOM and C-DOM are indicated in panels A and B. The TADkit tool was used to model Hi-C datasets from [34].

We applied the same 3D modeling viewer to Hi-C datasets obtained with limb bud cells from the *HoxD*^*Del(1-13)d9lac*^ mutant mouse stock (hereafter *Del(1-13)d9lac*), which contains a deletion including the *HoxD* cluster [34] (see Additional File 1). In this mutant, the deleted DNA was substituted by a *Hoxd9lac* reporter transgene while the *Evx2* and *Lnpk* genes remain present. In the absence of the *HoxD* cluster, C-DOM and T-DOM were still observed as independent structures despite a substantial shortening of the distance separating them (Additional File 1B-C). A clear spatial contraction was nevertheless scored between C-DOM and the first sub-TAD in T-DOM until region CS38-41 (Additional File 1B, C, red disk).

We next used the *HoxD*^*del(attP-Rel5)d9lac*^ (hereafter *Del(attP-Rel5)d9lac*) Hi-C datasets from mutant limbs lacking ca. 350kb of DNA including the *HoxD* cluster plus flanking regions (Fig. 1A, C). With this large deletion, the two TADs merged into one single structure (Fig. 1C) regardless of the cell type considered (distal or proximal), indicating that the TAD boundary had been entirely deleted. In this stock, the same *Hoxd9lac* transgene could be used as a transcriptional readout. The consolidation of T-DOM and C-DOM into one single structure was obvious up to the CS38-41 region, whereas the most telomeric located sub-TAD in T-DOM was somewhat less engulfed (Fig. 1C). We also computed an eigenvector analysis and distributed the interacting domains according to the first eigenvector values. We concluded that the position of the *HoxD* locus in compartment A, as well as the general compartment distribution along chromosome 2, were virtually identical between distal and proximal cells, when both the wild type and the *del(attP-Rel5)d9lac* datasets were considered (Additional File 1D).

### Transcription at the *HoxD* locus in the absence of the *HoxD* cluster

We looked at transcription emanating from the *lacZ* reporter transgenes by whole-mount *in situ* hybridization (WISH) on E11.5 fetuses, using a *LacZ* RNA probe and could identify both the distal and proximal limb domains in the two *del(attP-Rel5)d9lac* and *del(1-13)d9lac* lines, though with subtle variations in their relative strengths (arrowheads Additional File 2A). Therefore, even in the complete absence of a TAD boundary, the functional partition of proximal and distal enhancers occurred in a close-to-normal manner, with a clear separation between the two expression domains. While the distal domain overlapped well with the wild type *HoxD* distal limb pattern, the proximal domain was somewhat different in shape and position from the wild type *Hoxd9* pattern, resembling the expression pattern of the *Hog* lncRNA [34] thus likely indicating some enhancer reallocation due to the novel topology of the locus.

In order to have a complete account of such local modifications in transcriptional responses following the fusion of the two TADs, we carried out RNA-seq for both proximal and distal cell populations in control (*Wt*) and *del(attP-Rel5)d9lac* mutant limbs at E12.5. In control proximal cells, transcripts were expectedly detected both at the *HoxD* cluster, at the flanking *Lnpk* and *Mtx2* genes as well as for the *Hog* and *Tog* lncRNAs, two non-coding RNAs localized within T-DOM and normally responding to T-DOM enhancers [34, 37] (Fig. 2A, top). In control distal cells, while the expression of the latter two lncRNAs was undetectable, digit-specific transcripts were scored over the Island3 region both by RNA-seq and by WISH (Fig. 2B, top and Additional File 2B), a region previously defined as a distal cells-specific enhancer [38]. Therefore, we used these non-coding RNAs (*Hog, Tog* and Island3) as proxys to evaluate the activity of their surrounding proximal *versus* distal enhancers in the absence of the target *Hoxd* genes.

**Fig. 2.**
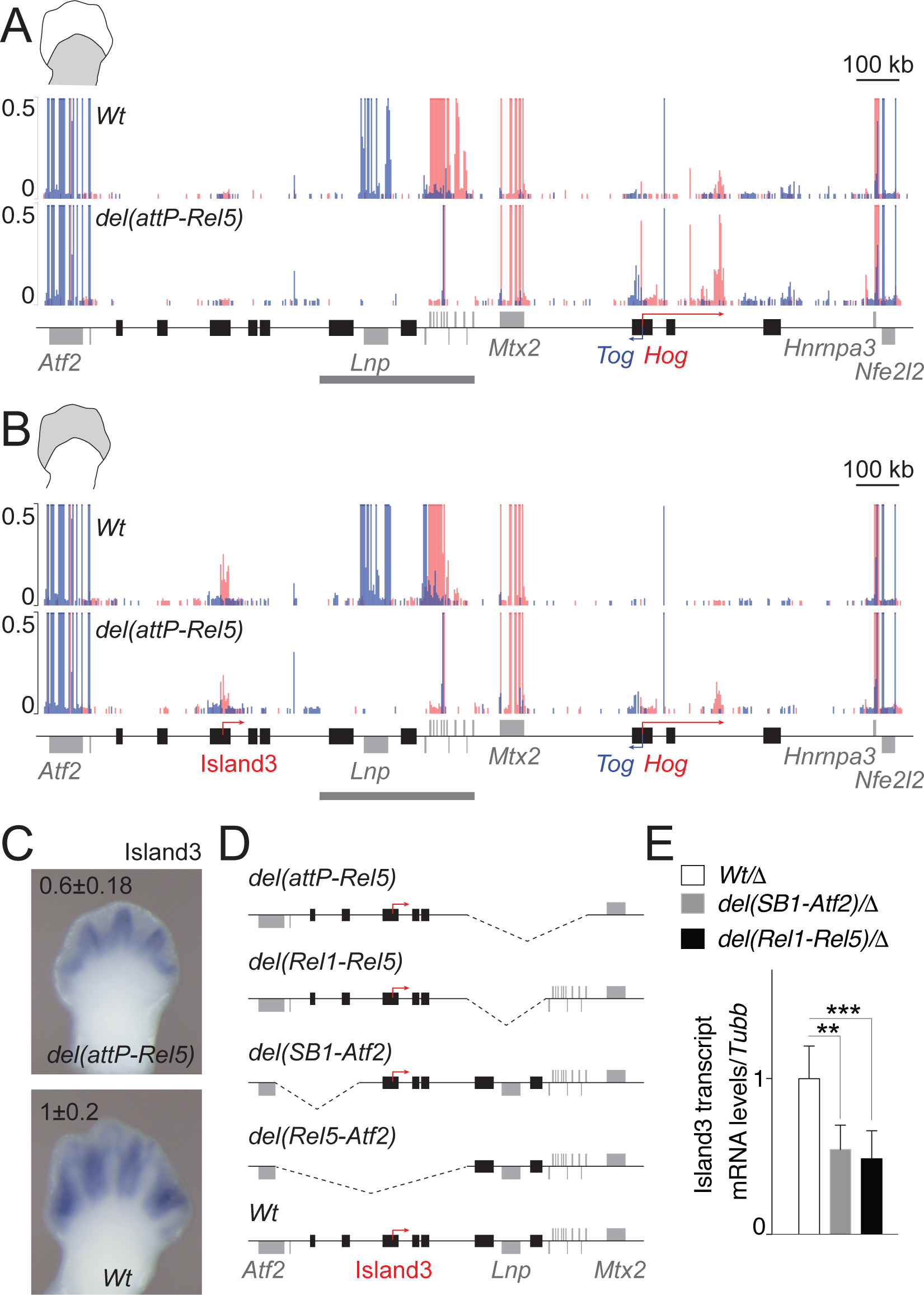
Transcript profiles at the *HoxD* locus in both control (*Wt*) and *del(attP-Rel5)d9lac* mutant limb buds. (*A, B*) Normalized RNA-seq profiles of control (*Wt*) and mutant proximal (A) and distal (B) limb cells. Values from forward (red) and reverse (blue) strands are merged into the same graph. The positions of various genes and of Island3 are shown below. The thick grey lines depict the *del(attP-Rel5)d9lac* deletion. The isolated signal around *Hoxd9* in the second tracks in *A* and *B* arises from the *Hoxd9/lacZ* reporter transgene present in the mutant line. The scale is set such that changes in non-coding regions can be better observed. *n*=3. (*C*) WISH of Island3 eRNA in both *del(attP-Rel5)d9lac* and wild type E12.5 forelimbs. qPCR values (mean±SD) are shown on the top of each image. *n*=6 for *Wt* and *n*=4 for *del(attP-Rel5)d9lac*. (*D*) Schemes of the various deleted regions of the mutant lines used in panels *A* to *E*. (*E*) qPCR of Island3 eRNAs in E12.5 distal limb cells in two distinct partial deletions of C-DOM. The mutant lines used were *del(SB1-Atf2)* (*n*=4) and *del(Rel1-Rel5)* (*n*=9), both balanced by the *del(Rel5-Atf2)* (*n*=12) allele (where Island3 is deleted, abbreviated by Δ in the legend). Results were compared to *del(Rel5-Atf2)/+* samples as controls (white bar). Bars show mean ± SD. Welch’s *t*-test ** p=0.0026 and *** p<0.0001.

In proximal mutant limb cells, the *Hog* and *Tog* RNA levels substantially increased (adjusted p-value from DESeq2 analysis of 1.75e-10 and 6.72e-22, respectively) while at the same time, the mRNA levels corresponding to the housekeeping genes *Mtx2* and *Atf2* remained approximately the same (adjusted p-value=1.00) (Fig. 2A, bottom and Additional File 2C). Transcripts for *Hoxd* genes and *Lnpk* had expectedly disappeared after the deletion, yet a signal remained for *Hoxd9* reflecting the transcription of the reporter gene left in place. Of note, the level of Island3 e-RNA did not seem to increase in the deleted configuration. Therefore, while in the absence of target *Hoxd* genes, proximal enhancers within former T-DOM were partly re-allocated towards the *Hog* and *Tog* promoters, they did not seem to affect Island3 expression, despite the removal of the TAD boundary (Fig. 2A, bottom, Additional File 2C).

In distal limb cells, the level of Island3 e-RNA decreased in the deleted configuration (Fig. 2B, C). While this transcript did not appear as differentially expressed in our RNA-seq whole genome analysis due to restrictive parameters (46% reduction mutant versus control, p-value=1.4e-4; Additional File 2D), it showed a significant reduction by qPCR (40% reduction mutant versus control, Welch’s *t*-test p-value= 0.0166) and by WISH (Fig. 2C). Likely, this decrease of expression was due to the loss of the GCR and Prox distal enhancers, as suggested by the deletion of the Rel1 to Rel5 region. A comparable outcome was observed in the deletion SB1 to *Atf2*, which removes two different enhancers (island1 and 2) on the other end of the regulatory domain (Fig. 2D, E). Noteworthy, neither of the housekeeping transcription units was transcribed more efficiently. However, a significant increase in *Hog* and *Tog* lncRNAs was scored, while these two genes are normally silent in distal cells where T-DOM has been switched off (Fig. 2B, Additional File 2D). Such an up-regulation could illustrate either a weakening in T-DOM repression in distal cells, or novel interactions between distal enhancers located in former C-DOM and the two lncRNAs’ loci, following the deletion of the TAD boundary.

### Changes in chromatin marks after TADs fusion

We complemented these observations by looking at the acetylation of H3K27, using proximal and distal E12.5 limb bud tissue derived from both control and *del(attP-Rel5)d9lac* fetuses. In proximal cells, the distribution of H3K27ac marks in the mutant material was as in control (wild type) cells (Fig. 3A). H3K27ac modifications were found enriched in T-DOM (the active TAD) while depleted from C-DOM (the inactive TAD). The amount of H3K27ac was slightly increased over a large region of T-DOM in mutant cells, with a particular increase at the transcription start site of both *Hog* and *Tog* (Fig. 3A, 120% increase, arrowhead), thus matching the previously described increased in RNA levels (Fig. 2). The distribution of H3K27ac marks over C-DOM was comparable in control and mutant proximal cells (Fig. 3A, see *del versus Wt*).

**Fig. 3.**
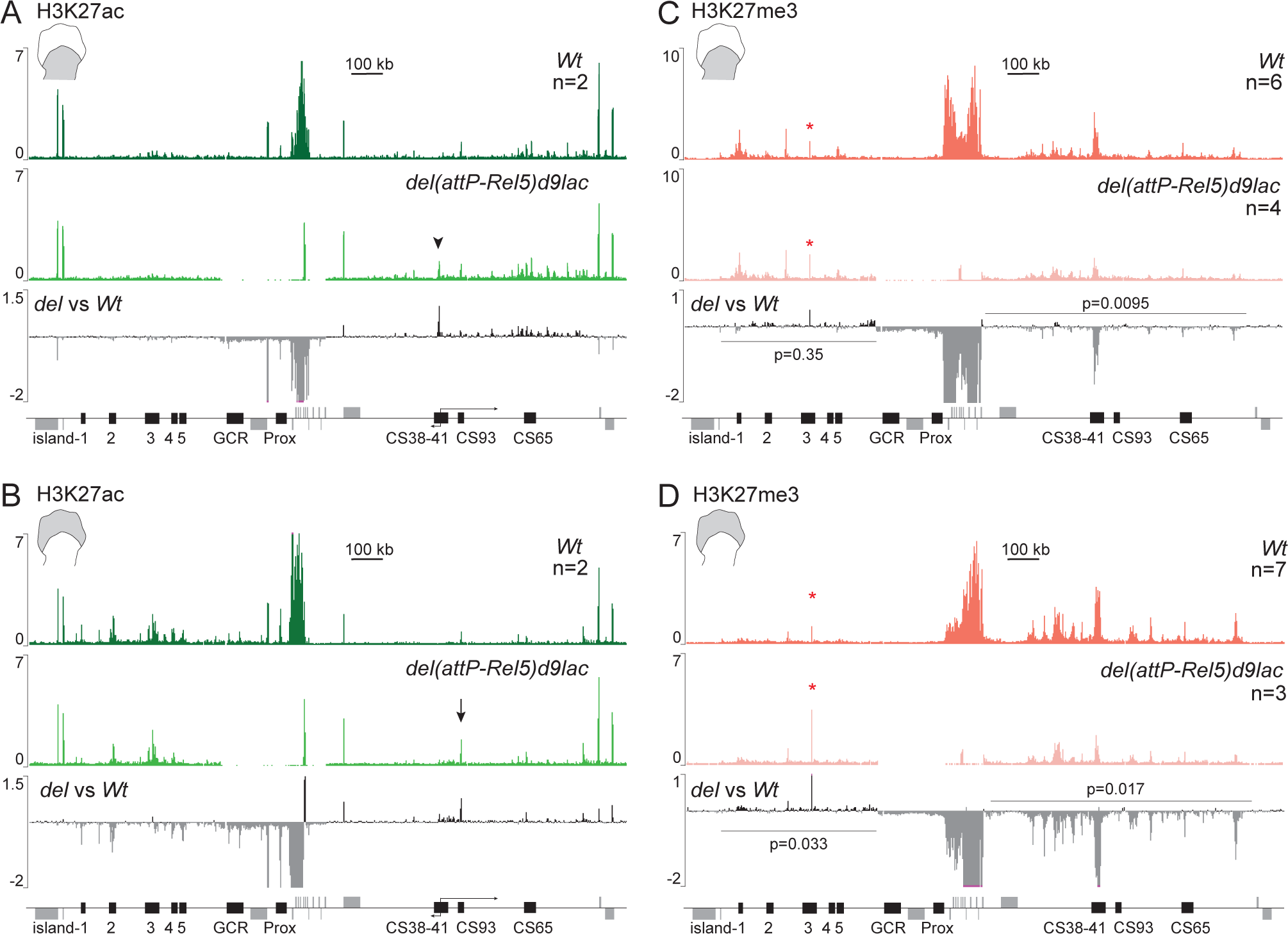
Distribution of H3K27ac (A, B) and H3K27me3 (C, D) marks over the *HoxD* cluster and its flanking TADs in both control (*Wt*) and *del(attP-Rel5)d9lac* proximal (A, C) and distal (B, D) limb bud cells. *(A, B*) H3K27ac ChIP profiles from proximal (A) and distal (B) limb cells. Control is on top and the *del(attP-Rel5)d9lac* profile is shown below along with the difference of deleted versus control ChIP data (*del vs Wt*). The arrowhead in A depicts the shared *Hog* and *Tog* start site (see also the divergent arrows below). The arrow in B indicates the CS93 enhancer. (*C, D*) H3K27me3 ChIP profiles from proximal (C) and distal (D) limb bud cells. Control is on top and the *del(attP-Rel5)d9lac* track is shown below along with a comparison profile showing the difference between mutant and wild-type profiles. The data were averaged between different experiments (*n* is shown on the right). The red asterisks indicate artifactual peaks. The signal from the *Hoxd9* region in the deleted allele corresponds to the *Hoxd9/lacZ* transgene.

In *del(attP-Rel5)d9lac* mutant distal cells, an increase in H3K27ac was scored at region CS38-41 (Fig. 3B, 75% increase), which correlated with the activation of these two lncRNAs in these mutant cells, while they are normally silent in their wild-type counterparts (Fig. 2B). Additionally, a strong increase in this histone mark was scored in CS93 (Fig. 3B, arrow, 75% increase), a region recently characterized as a proximal limb enhancer [15]. The general distribution of H3K27ac appeared slightly increased throughout T-DOM in mutant cells when compared to control (Fig. 3B). This slight increase in T-DOM activity was also noticeable when analyzing proximal mutant tissue. A striking effect was however observed in H3K27ac coverage over C-DOM, in mutant *versus* control distal cells. A substantial loss of H3K27ac was indeed scored over the regulatory regions island 1, 2, 4 and 5 (Fig. 3B, about 40% decrease). This effect was not as evident over island 3, i.e. in the region where the enhancer transcript was detected in both control and mutant distal cells (Fig. 2B). Therefore, in distal cells, the fusion of both TADs and removal of target genes seemed to weaken the transcriptional activity of C-DOM, while maintaining the activity of T-DOM well above the silencing observed in control distal cells.

To further document this observation, we looked at the distribution of H3K27me3 marks. In control proximal limb cells, H3K27me3 were detected over T-DOM at E12.5 (Fig. 3C), i.e. when this landscape is still functionally active, likely due to the presence of a large percentage of negative cells in the dissected material (see [33]). In distal cells, where T-DOM is switched off, a robust increase was detected with a strong coverage of the entire T-DOM (Fig. 3D). In proximal cells, H3K27me3 marks were also scored over the silent C-DOM regulatory islands, a labeling that mostly disappeared upon the activation of these regulatory islands in distal cells (Fig. 3D). The H3K27me3 profiles obtained with the *del(attP-Rel5)d9lac* mutant limb buds were in agreement with the distributions of both the H3K27ac marks and the transcripts. In proximal mutant cells, the profile was globally similar to that seen in control cells with however a 50 percent decrease at region CS38-41 (Fig. 3C). In distal cells, the same effect was scored, yet at a much higher magnitude. H3K27me3 marks were heavily depleted from T-DOM whereas they were found mildly but significantly enriched all over the C-DOM region containing the regulatory islands (Fig. 3D, respectively 50% decrease and 20% increase). Therefore, these results confirmed that in mutant cells carrying the combined neoTAD, the former T-DOM landscape is globally overactive in distal cells, at the expense of C-DOM enhancers, which appear less active than in their native context.

### Recruitment of PRC complexes at the *HoxD* cluster and surroundings

Polycomb repressive complexes (PRC1 and PRC2) are generally associated with lack of gene expression and usually recruited to CpG islands close to transcriptionally active regions [24, 39, 40]. In this context, the massive presence of H3K27me3 marks over T-DOM, a region largely devoid of coding units, raised the question of the recruitment mechanism at work. We looked at the presence of both EZH2 and RING1B, two components of PRC2 and PRC1, respectively. ChIP experiments revealed that EZH2 was located mostly within the *HoxD* cluster (Fig. 4A). Outside the gene cluster, a weak enrichment was scored over region CS38-41 in proximal cells, which appeared even weaker in distal cells. Altogether, the two gene deserts were generally devoid of PRC2. A comparable conclusion was reached regarding the prevalence of signal at the cluster, with the analysis of the PRC1 component RING1B, even though some enrichment was detected on the gene deserts, generally over T-DOM and particularly over the CS38-41 and CS65 regions, without any striking difference between distal and proximal cells (Fig. 4A). Some light differences were scored in C-DOM, where a few regulatory regions appeared specifically decorated in proximal tissue but devoid of RING1B in distal tissue (compare Island1 and Island4 in Fig. 4A).

**Fig. 4.**
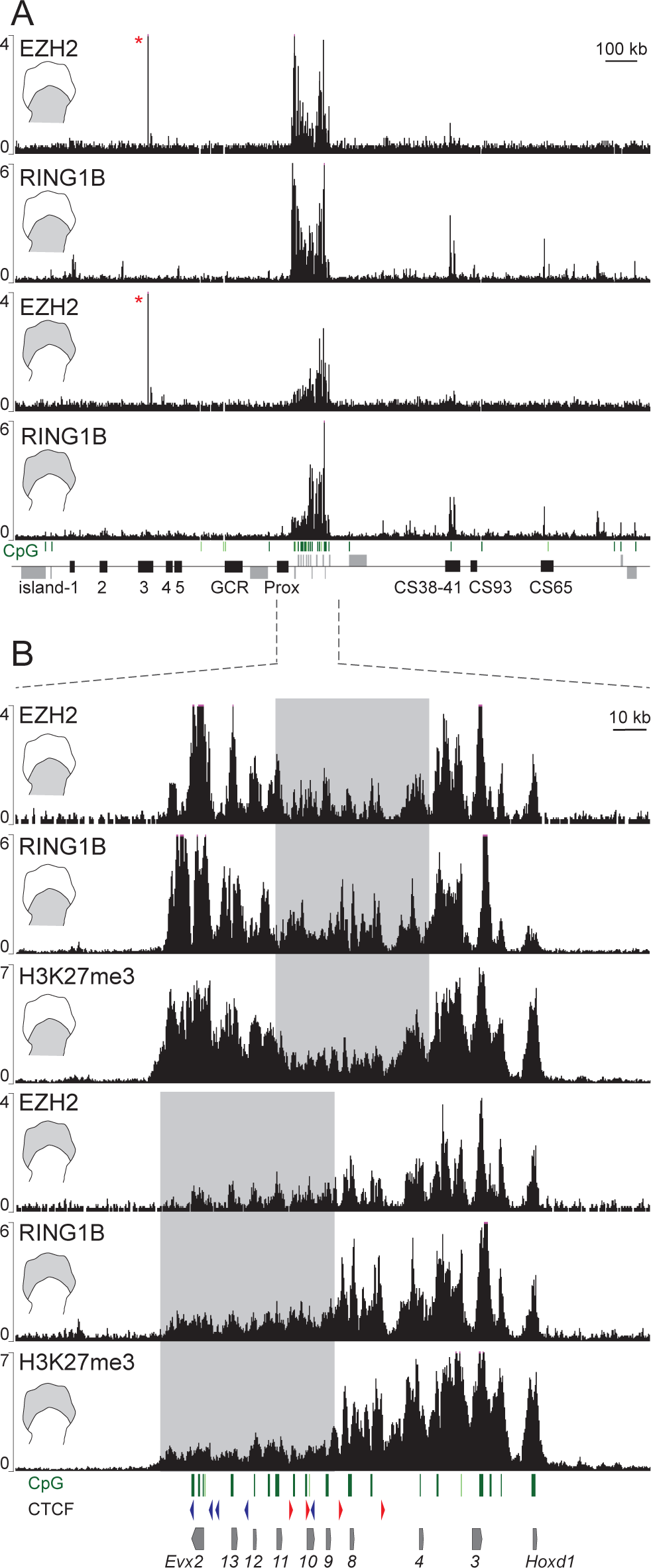
Distribution of PRC1 (RING1B) and PRC2 (EZH2) over the *HoxD* cluster and regulatory landscapes in limb bud cells. (*A*) EZH2 and RING1B ChIP profiles in proximal (top two panels) and distal (bottom two panels) E12.5 limb bud cells. The CpG distribution is shown as green bars on top of the gene diagram. The red asterisk indicates an artifactual signal. (*B*) Magnification of the *HoxD* cluster showing the distribution of EZH2 and RING1B in proximal and distal limb cells. H3K27me3 ChIP tracks are shown for each tissue. The CpG islands are shown as green bars and the CTCF and their orientations are depicted as blue (reverse strand) or red (forward strand) arrowheads.

Within the *HoxD* cluster itself, the distribution of both EZH2 and RING1B nicely matched the coverage by H3K27me3 in its general and tissue-specific extents (Fig. 4B) <u>[1–3]</u>. In proximal cells, the coverage was minimal over those genes active in response to T-DOM enhancers (from *Hoxd8* to *Hoxd11*, rectangle in Fig. 4B, tracks 1 and 2), while in distal cells genes responding to C-DOM enhancers were bound only weakly by either PRC2 or PRC1 (from *Hoxd13* to *Hoxd10*, Fig. 4B, rectangle in tracks 4 and 5). The EZH2 signals were significantly enriched at CpG islands and over coding regions, whereas the distribution of PRC1 was broader (Fig. 4B), suggesting a recruitment of PRC2 by CpG islands [24, 40, 41].

Considering that H3K27me3 covered both *Hox* genes and their regulatory landscapes, whereas PRC complexes were mostly recruited to the *HoxD* cluster itself, we wondered whether the reduction of H3K27me3 marks along T-DOM in *del(attP-Rel5)d9lac* mutant proximal cells could result from the mere absence of the *HoxD* gene cluster. To this aim, we used the engineered *HoxD*^*inv(attP-Itga6)*^ inversion (hereafter *inv(attP-Itga6)*), where the *HoxD* cluster was disconnected from T-DOM and displaced circa 3Mb away while preserving both its integrity and its association with C-DOM [42] (Additional File 3).

We verified that the genomic interactions between *Hoxd* genes and T-DOM were abrogated in this *inv(attP-Itga6)* inverted allele by performing a 4C-seq analysis in mutant and control distal limb cells, with *Hoxd4* and CS38 as viewpoints (Fig. 5A). Expectedly, the contacts established by *Hoxd4* were no longer oriented towards T-DOM in the inversion allele, when compared with control (Fig. 5A, tracks 1 and 2). In this inverted allele, interactions were now established *de novo* between *Hoxd4* and a region around the *Itga6* and *Dlx1/Dlx2* genes, near the inversion breakpoint. Also, contacts with C-DOM were slightly increased. Furthermore, when region CS38 was used as a viewpoint, interactions with the *HoxD* cluster were largely lost and most contacts remained within T-DOM itself (Fig. 5A, tracks 3 and 4).

**Fig. 5.**
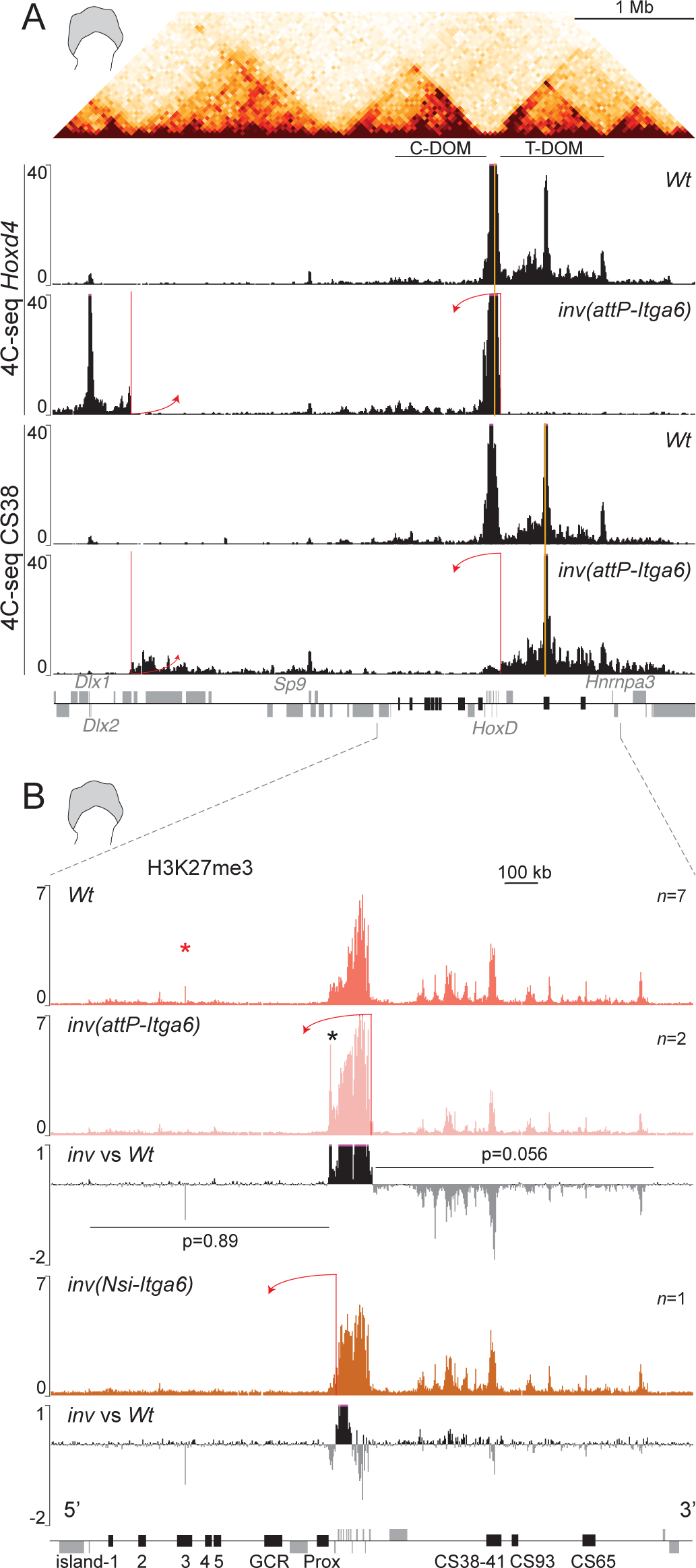
Epigenetic changes after disconnecting the *HoxD* cluster from its flanking T-DOM. (*A*) On top, a Hi-C profile of distal limb bud cells shows the *HoxD*-associated TADs. The panels below show a comparison of 4C-seq tracks between control (*Wt* from [34]) and *inv(attP-Itga6)* mutant distal limb cells. Either the *Hoxd4* gene (top two panels), or the CS38 region (bottom two panels), was used as baits (yellow vertical bars). The red bars indicate the locations of the loxP sequences used to generate the inversion. After inversion, contacts between *Hoxd4* and T-DOM are all lost, while they barely change when region CS38 is used as bait. (*B*) H3K27me3 ChIP profiles in control (*Wt*) and either the *inv(attP-Itga6)* inversion (top two profiles), or the *inv(Nsi-Itga6)* inversion (bottom two profiles). Below each mutant track, a comparison between mutant and control data is shown. The red bars indicate the inversion break points. In the *inv(attP-Itga6)* track, an additional peak appears at the 5’ extreme of the *HoxD* cluster (black asterisk), corresponding to an ectopic sequence introduced when building the attP breakpoint. The red asterisk indicates an artifactual signal. The number of replicates is shown for each track. Below each mutant track, a difference profile of mutant *versus* control signals is represented.

In this inverted configuration, the global amount of H3K27me3 marks deposited over T-DOM was substantially lower when compared to the control cells (Fig. 5B, tracks 1 and 2). This decrease was not observed when another inversion was used as a control. In the *HoxD*^*inv(Nsi-Itga6)*^ allele (hereafter *inv(Nsi-Itga6)* [43] the *HoxD* cluster remains in place yet C-DOM is inverted towards the same *Itga6* breakpoint. Therefore, these two inversions are identical except that one contains the *HoxD* cluster whereas the other does not (Fig. 5B, arrows in tracks 2 and 4 and additional File 3). In the *inv(Nsi-Itga6)* inversion allele, the enrichment of H3K27me3 over T-DOM was not decreased, as was the case for the *inv(attP-Itga6)* allele (Fig. 5B), neither in distal cells, nor in proximal cells (Additional File 3B). Altogether these results and those obtained with the *del(attP-Rel5)d9lac* allele, suggest that the presence of *Hoxd* genes was necessary to achieve a full spread of H3K27me3 marks over T-DOM, up to 800kb in far*-cis*.

Interestingly, this effect was restricted to T-DOM, as seen after zooming out and looking at a 10 Mb interval surrounding the *HoxD* cluster. In control distal cells, the distribution of H3K27me3 marks was enriched selectively over T-DOM, terminating abruptly at its TAD boundary with no further telomeric spreading. In mutant *del(attP-Rel5)d9lac* distal cells, despite the large reduction of H3K27me3 signals, the remaining coverage was also restricted up to the new telomeric boundary of the neoTAD (Additional file 4A). Comparable results were obtained when comparing the mutant *inv(attP-Itga6)*. In all cases, though to a different extent, the TAD structure appeared to determine the extent of H3K27me3 spreading.

### H3K27me3 inheritance and clearance

The *inv(attP-Itga6)* allele disconnected T-DOM proximal enhancers from their target *Hoxd3 to Hoxd11* genes, similar to a previous case when a deletion of T-DOM was used [33]. In both cases, expression of these target genes was expectedly lost in proximal cells of the forelimb buds (Fig. 6A, B; see also [33]). Unexpectedly however, both the quantity and distribution of *Hoxd9* and *Hoxd11* mRNAs (see digits II and V) were also reduced in distal cells, where these genes are under the control of C-DOM enhancers (Fig. 6A, B, arrows and arrowheads respectively). This surprising observation was explained by the lineage transmission, from proximal to distal cells, of H3K27me3 marks abnormally present in *Hoxd* genes in the absence of T-DOM [33].

**Fig. 6.**
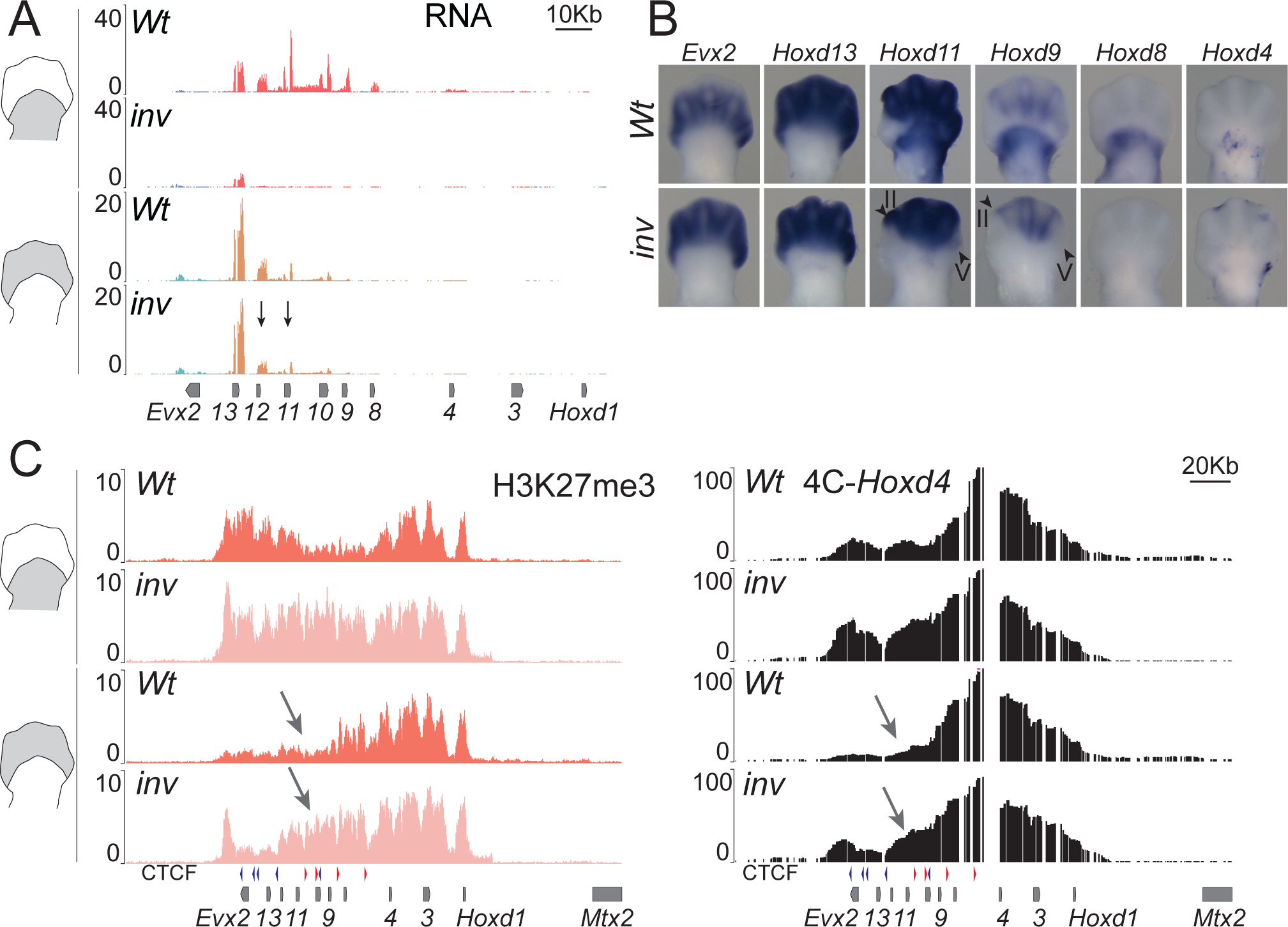
*Hoxd* gene expression in limbs in the absence of T-DOM. (*A*) Normalized RNA-seq profiles of control (*Wt*) and *inv(attP-Itga6)* mutant proximal (A) or distal (B) limb bud cells. Black arrows indicate the decreased RNA quantity over *Hoxd12* and *Hoxd11* in distal tissue (bottom two tracks) while expression has almost fully disappeared in proximal limb cells (top two tracks). (*B*) WISH of *Hoxd4, Hoxd8, Hoxd9, Hoxd11, Hoxd13* and *Evx2* in E12.5 forelimb buds. The arrowheads indicate digits II and V. (*C*) On the left, comparison of H3K27me3 signal over the *HoxD* cluster in either proximal (top two tracks) or distal (bottom two tracks) between control (*Wt*) and mutant *inv(attP-Itga6)* specimen. The CTCF sites are shown below. The arrows point to the extension of the H3K27me3 negative domain over the *Hoxd11* region in mutant *inv(attP-Itga6)* distal cells (fourth track), when compared to control cells (third track). In the right, 4C-seq tracks showing interactions inside the *HoxD* cluster when *Hoxd4* is used as bait (*Wt*: data from [34]). The arrows indicate a robust gain of interaction over *Hoxd11* to *Hoxd12* region in *inv(attP-Itga6)* mutant distal cells.

To further assess this possibility, we analyzed the precise distribution of H3K27me3 marks over the *HoxD* cluster in the *inv(attP-Itga6)* allele. In proximal cells, we found a high and homogeneous coverage of this histone modification, from *Hoxd1* up to *Evx2*, unlike in the control allele where the DNA interval between *Hoxd8* and *Hoxd11* was transcriptionally active and hence depleted from this mark (Fig. 6C, tracks 1 and 2). Accordingly, the homogeneous distribution of H3K27me3 over the gene cluster in the mutant allele reflected the complete lack of *Hoxd* expression in proximal cells (Fig. 6A tracks 1 and 2 and Fig. 6B). In control distal cells, the region from *Evx2* to *Hoxd9* was depleted in H3K27me3 marks, as expected from the regulation of C-DOM enhancers. In inverted mutant cells however, an abnormally high H3K27me3 coverage was scored over the *Hoxd9* to *Hoxd11* region (Fig. 6C, arrow in track 4), which corresponded to the decrease in transcript levels observed for these genes in mutant distal cells (Fig. 6A tracks 3 and 4). This increase in H3K27me3 was not observed in the *inv(Nsi-Itga6)*, where these genes are normally expressed in proximal tissue (Additional File 5). Because distal limb bud cells are the descendants in lineage of proximal cells (see [44], we explain this negative effect over C-DOM regulation by the transmission of H3K27me3 to distal cells. This mark was ectopically detected over the *Hoxd4* to *Hoxd11* region in proximal cells, due to the lack of contacts between proximal enhancers and their target *Hoxd* genes, thus preventing their transcriptional activation. Of note, *Hoxd13* and *Evx2* transcript levels remained unchanged in the mutant allele when compared to control.

We assessed whether this ectopic gain of H3K27me3 in proximal cells would translate into a change in the extent of the negative chromatin sub-domain formed at *Hox* loci by H3K27me3-enriched sequences [45, 46]. We carried out 4C-seq by using *Hoxd4* as a viewpoint and noticed that in proximal cells, contacts established by *Hoxd4* clearly extended over the centromeric part of the cluster in the mutant allele, in agreement with the gain of H3K27me3. These contacts were also observed, though to a slightly lesser extent, in mutant distal cells, again correlating with the persistence of H3K27me3 marks (Fig. 6C, arrow in track 4).

## DISCUSSION

During limb development, the two TADs associated with the *HoxD* cluster are either transcriptionally active or repressed, in an exclusive manner. Initially, T-DOM enhancers are active and control the first wave of *Hoxd* transcription in early limb buds and, subsequently, in proximal structures such as the forearms [33]. In a second phase, C-DOM enhancers become activated in distal limb (the future hands and feet) while T-DOM concomitantly terminates operating and becomes covered by negative H3K27me3 marks [33, 38]. This bimodal regulation in TAD activities is necessary to organize each of the proximal and distal *Hox* expression domains, which are essential for proper limb development [47–50].

### A fused neoTAD

Studies addressing the mechanism underlying the functional switch between these two TADs have suggested that in this particular case, TADs could represent coherent and independent regulatory units, i.e. that the 3D structure itself may participate to the global functional output of the system. In this view, a TAD could be either functionally permissive or refractory to the implementation of all the enhancers it may contain [32] thus representing an additional regulatory layer. In the case of T-DOM and C-DOM, only one of them is licensed to work at a time since the presence of HOX13 proteins, partly determined through the activation of C-DOM, leads to the repression of T-DOM [32]. We thus wondered how this functional exclusivity would translate after the fusion of the two structures, in a situation where both proximal and distal enhancers would be included in the same neoTAD. In this neoTAD indeed, several enhancers normally present in C-DOM, i.e. with a distal specificity, were now located along with enhancers normally displaying a proximal specificity due to their location within T-DOM. Since their genuine target genes (*Hoxd*) were absent, we assessed their functionality by using three transcription units as readouts: an eRNA encoded by Island3 within former C-DOM, the *Hog* and *Tog* lncRNAs encoded within former T-DOM and a *Hoxd9/lacZ* reporter transgene positioned exactly between the former two TADs.

The analysis of *lacZ* mRNA revealed the presence of distinct proximal and distal expression domains, suggesting that the presence of the two kinds of enhancers in the same neoTAD did not drastically affect neither their global functional specificities, nor their mode and sequence of action. However, the proximal domain was distinct from what is normally observed in wild type limbs, despite the remaining presence of all known proximal enhancers in the two deleted alleles. In fact, it resembled in its position and shape to the expression domain of the lncRNA *Hog*, which lies in the vicinity of proximal enhancers within T-DOM. In this case, the absence of target genes and their associated CTCF sites may have led reallocations in enhancer-promoters contacts, as also suggested by the upregulation of *Hog* and *Tog* lncRNAs in proximal mutant cells. Therefore, the final transcription readout of T-DOM enhancers may slightly vary in space and time depending on how the target promoters are organized and on their local topology. Furthermore, *Hog* and *Tog* transcripts were scored in mutant distal cells, while completely switched off in control distal cells. We interpret this as a response to the remaining C-DOM enhancers, in the absence of the TAD boundary. Also, the repression of T-DOM in mutant distal cells was not implemented as efficiently as in control cells, thus contributing to this light up-regulation. Housekeeping genes located within or in the vicinity of former T-DOM, such as *Mtx2, Hnrnap3* or *Atf2*, were left transcriptionally unaffected after fusion of the TADs, as these genes were not able to respond to the liberated enhancers.

In parallel with the maintenance, in the neoTAD, of proximal enhancer activity in distal cells, the level of Island3 eRNA was slightly reduced. While this RNA was present in control distal cells but absent from control proximal cells, the same regulatory region, after TAD fusion, showed a diminution of its transcriptional activity, as if the neoTAD was globally pushed towards a proximal type of regulation. A clear distal domain was nevertheless detected with the *lacZ* expression pattern, demonstrating the activity of at least some distal limb enhancers and hence the reduction in Island3 eRNA may also be caused by the deletion of some distal enhancers in the neoTAD.

This tendency of the neoTAD to adopt a type of regulation, which generally speaking appeared more proximal than distal, was reinforced by the analyses of chromatin marks. In distal cells, the fusion between the two TADs was indeed accompanied by a decrease in H3K27ac coverage in several enhancers located in former C-DOM. In contrast, H3K27ac marks in mutant distal cells were more abundant in the former T-DOM region, i.e. over proximal enhancers, than in control distal cells where these marks rapidly disappear [33]. In general terms, however, H3K27ac deposition associated to enhancer activation in mutant cells was still observed as in control cells, indicating that former T-DOM enhancers were still active in proximal limb bud cells and former C-DOM enhancers in distal cells. The difference was observed in the balance between these two types of regulations, rather than in their implementation.

The profile of H3K27me3 marks again confirmed these observations. In distal cells, i.e. in cells where T-DOM is normally inactive and hence most of the TAD is decorated with such marks, the amount of H3K27me3 was significantly reduced in mutant *versus* control cells, as if the ‘proximal regulation’ had not been entirely switched off, even in distal cells. In parallel with both the decrease of Island3 eRNAs and the decrease in H3K27ac, the distribution of H3K27me3 marks appeared increased in the former C-DOM region.

Altogether, these results suggest that when mixed into a single neoTAD, the proximal regulation tends to take the lead over the distal regulation, with proximal enhancers that are active for too long, even in distal cells where distal limb enhancers seem to be somewhat under-active. A potential mechanism may involve the reported effect of HOX13 proteins in the termination of T-DOM regulation, combined with the novel chromatin architecture of the neoTAD. In the absence of HOXD13 proteins, deleted from the neoTAD, the dose of HOXA13 should be sufficient to secure the repression of T-DOM and thus to implement the switch in regulations [32]. However, the new chromatin configuration of this part of T-DOM when included in the neoTAD may affect the negative function of HOXA13, leading to a partial inhibition only and hence to an improper switch off of proximal enhancers.

Nonetheless, this leakage in the strict regulatory switch observed at this locus under normal conditions does not prevent the two large proximal and distal expression domains to form, with the negative cellular domain in between, which is the landmark of a correct bimodal regulation at the *HoxD* locus. One possibility to consider is that in the neoTAD, the CS38-41 region induced an internal boundary between a large and novel chromatin domain containing parts of both C-DOM and T-DOM on the one hand, and the same telomeric sub-TAD in the wild type configuration, which contains most proximal enhancers as judged from chromatin modifications and 4C contacts profiles [15, 33]. Therefore, it is possible that this intra-TAD boundary allows for some isolation between proximal and distal enhancers to be conserved in the deletion mutant, as seems to be the case in the control situation. Further deletion of this boundary region including the CTCF sites should be indicative in this respect.

### TAD-specific and long-range effect of PRC silencing

Our results also provide some indications as to how PRC silencing propagates either *in-cis* at a distance or through cell divisions (see [51]). Within the *HoxD* cluster itself, we show that PRC2 recruitment selectively occurs at the CpG islands, as previously proposed (e.g. [24, 25, 52]. In addition, however, H3K27me3 marks were found throughout the T-DOM (over ca. 800Kb) in distal cells, where proximal enhancers have terminated their function, even though H3K27me3 marks were shown not to spread outside the *HoxD* cluster in a linear manner [53]. In the *del(attP-Rel5)d9lac* deletion, in the almost complete absence of CpG islands in and around the *HoxD* cluster, the enrichment by H3K27me3 marks in T-DOM was severely reduced in distal cells, indicating that indeed the recruitment of PRC2 complexes over the *HoxD* cluster was mandatory to start covering the telomeric regulatory landscape by H3K27me3 marks, concomitantly to its functional inactivation. Some H3K27me3 coverage was nevertheless detected in C-DOM and more substantially in T-DOM, perhaps due to the presence of both the *Hoxd9/lacZ* reporter transgene and the *Hog* and *Tog* transcription start sites.

Of note, the coverage by H3K27me3 marks in control distal cells outside the *HoxD* cluster itself, i.e. in a region that is not particularly enriched in PRC2, exactly matched the extent of the TAD containing those *Hoxd* genes inactivated in distal cells and hence heavily covered by PRC2, PRC1 and H3K27me3 (T-DOM). Such an effect was not scored in any other region in the 10Mb surrounding the *HoxD* locus. This result suggests that the global inactivation of T-DOM regulation in distal cells [32] is accompanied by a TAD-specific coverage of H3K27me3 marks, up to the telomeric TAD boundary where the presence of these negative marks abruptly stops (see also [54, 55]. Therefore, the TAD structure itself may dictate the extent of coverage by H3K27me3 marks, after recruitment of PRC2 by those *Hoxd* genes switched off in these distal cells and included into this TAD. In this view, the TAD (T-DOM) may be seen as a global functional unit.

### Heritability of Polycomb-associated gene silencing

During the replication of H3K27me3-labeled DNA sequences, daughter cells inherit this histone modification from their parental cell [51, 56, 57] Since limb development occurs mainly through a distal outgrowth, most distal cells, i.e. those where C-DOM regulation is at work, derive from proximal cells that used to be under the control of T-DOM enhancers. In the latter cells, the central part of the *HoxD* cluster is active and hence *Hoxd9, Hoxd10* and *Hoxd11* are devoid of H3K27me3 marks, whereas *Hoxd12* and *Hoxd13*, which are located on the other side of the TAD boundary are silent and thus covered by H3K27me3 marks [33].

When these cells become distal and start to implement the C-DOM regulation, H3K27me3 marks are erased from both *Hoxd13* and *Hoxd12*, the major targets of C-DOM enhancers, which are transcribed at high levels. Because *Hoxd11* and *Hoxd9* are devoid of H3K27me3 marks, they also become transcribed in distal cells, even though their genuine function in these cells has not been unequivocally demonstrated [37]. In the absence of T-DOM regulation in *inv(attP-Itga6)* proximal mutant cells, the entire *HoxD* cluster is heavily covered by H3K27me3 marks since all *Hoxd* genes are silenced. When these mutant distal cells start to implement the C-DOM regulation, the H3K27me3 marks covering *Hoxd13* and *Hoxd12* are removed with the same kinetics as in wild type distal cells, due to a comparable transcriptional context. However, *Hoxd11* and *Hoxd10* transcription onset is severely delayed when compared to control distal cells, as these genes were inherited in a silenced state, covered by H3K27me3 marks [33]. In this latter case, the strength of distal limb enhancers and the proximity of *Hoxd13* and *Hoxd12* likely leads to a progressive removal of PRC silencing and a weak and delayed activation of both *Hoxd11* and *Hoxd10* in distal cells. This observation illustrates both the capacity for cells to memorize their coverage in H3K27me3 marks in a physiological context, and the labile aspect of Polycomb silencing, which can be efficiently removed through a strong transcriptional activation.

## CONCLUSIONS

From this study, we conclude that proximal and distal limb enhancers, which are normally segregated between the two TADs flanking the *HoxD* cluster, were not dramatically affected neither in their activation, nor in their specificity, when their target genes were deleted and the two TADs merged into a single chromatin interaction domain. However, the modification in chromatin architecture occurring after the fusion of the two TADs affected the silencing of some enhancers and extended their activity over time. These results also suggest a mechanism whereby enhancer silencing is accompanied by a far-*cis* action of Polycomb group proteins after being recruited for the most part at target genes. Lastly, we conclude that active genes are more readily amenable to a subsequent enhancer regulation compared to silenced genes, illustrating the potential importance of Polycomb associated chromatin marks in the proper timing of gene activation during developmental.

## ADDITIONAL FILES

Additional_File_1.pdf

Additional_File_2.pdf

Additional_File_3.pdf

Additional_File_4.pdf

**Additional File 1.**
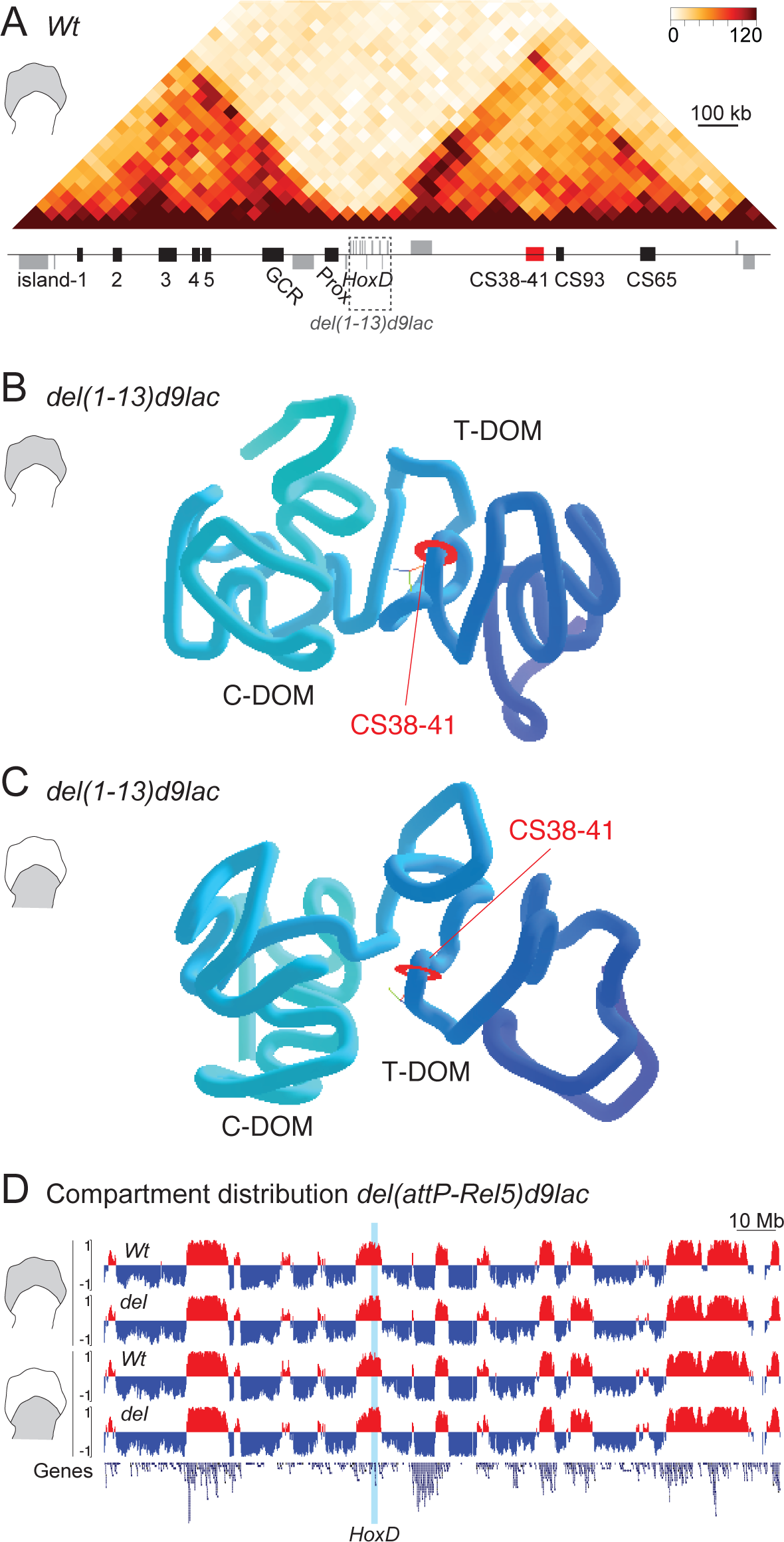
3D-representation of the *HoxD* locus in control (*Wt*) and *del(1-13)d9lac* mutant limb buds. (*A*) Hi-C map showing the presence of both TADs on either side of the *HoxD* locus in distal limb cells and its associated genes (grey boxes) and regulatory regions (black and red boxes). The dashed rectangle illustrates the deletion in the *del(1-13)d9lac* allele. (*B, C*) TADkit-derived 3D representation of Hi-C datasets [34] obtained for distal (B) and proximal (C) limb cells processed from *del(1-13)d9lac* mutant mice. The CS38-41 region is shown as a red disk in the 3D models to be used as a reference point. In this deletion allele, both TADs are still visible unlike in the larger *del(attP-Rel5)d9lac* deletion shown in Fig. 1. (*D*) A/B compartment distribution along chromosome 2. Eigenvectors were calculated from *Wt* and *del(attP-Rel5)d9lac* E12.5 distal and proximal limb Hi-C data. Compartment A is represented as positive values (red) and compartment B as negative values (blue). Gene density is shown in the bottom panel and the *HoxD* locus is indicated as a blue bar.

**Additional File 2.**
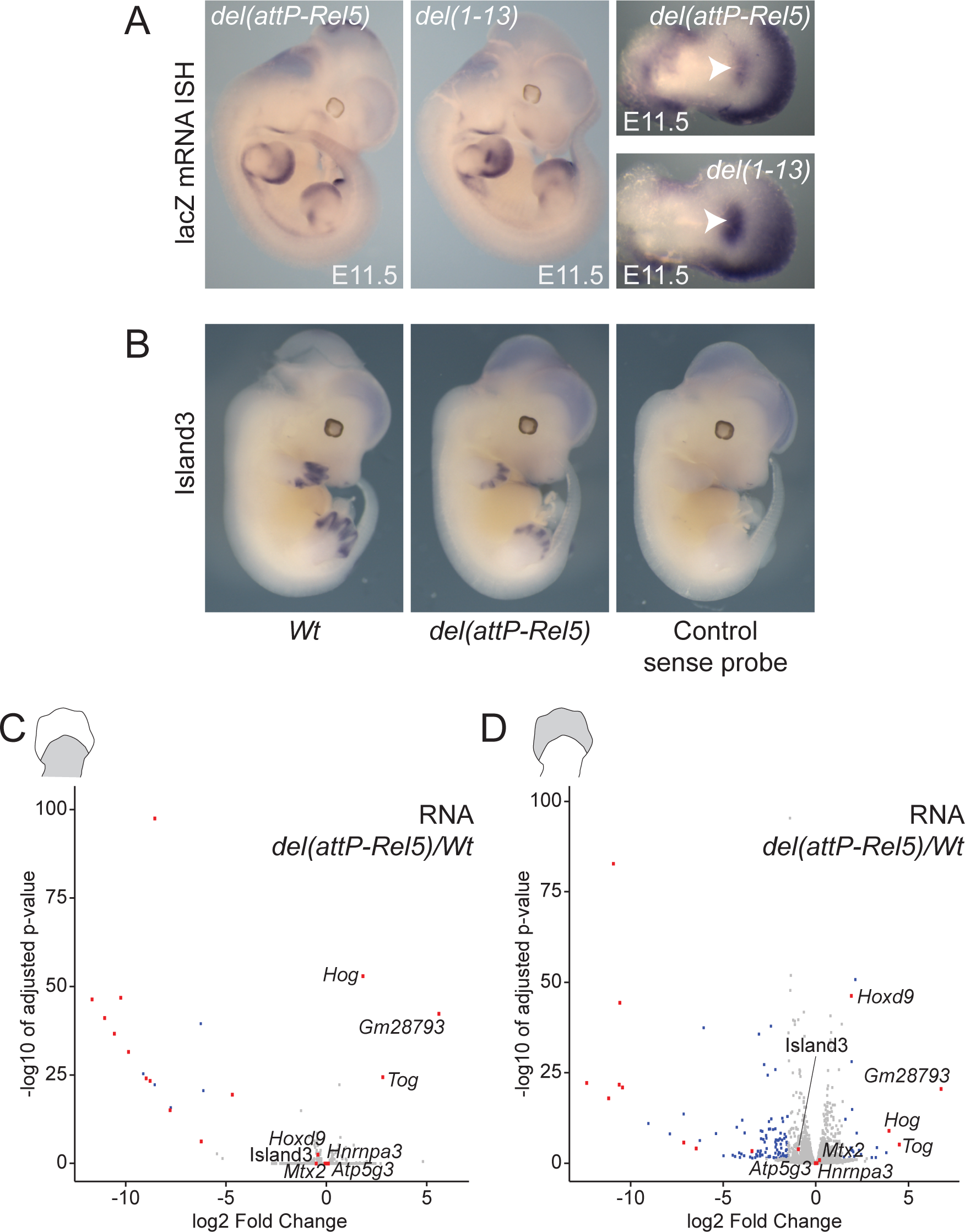
Expression analysis around the *HoxD* associated transcripts. (*A*) WISH of *lacZ* mRNA in *del(attP-Rel5)d9lac* and *del(1-13)d9lac* E11.5 forelimbs (right) and whole embryos (left). The proximal domain is shown by an arrowhead. (*B*) WISH of Island3 eRNAs in *Wt* and *del(attP-Rel5)d9lac* E12.5 embryos where the antisense probe was used (left) showing the specificity on the digital region. On the right, a probe control shows the lack of staining with the Island3 sense probe. (*C, D*) Volcano plots of all genes analyzed by RNA-seq in proximal (C) and distal (D) limb tissues comparing *del(attP-Rel5)d9lac* expression values to the control. All the genes located inside or in the vicinity of C-DOM and T-DOM are marked in red. Blue dots represent differentially expressed genes (absolute log2 fold change above 1.5 and adjusted p-value below 0.05) that are located outside these regions. *Hoxd9* and *Gm28793* (a short antisense mRNA) are significantly expressed due to their presence inside the *Hoxd9/LacZ* transgene.

**Additional File 3.**
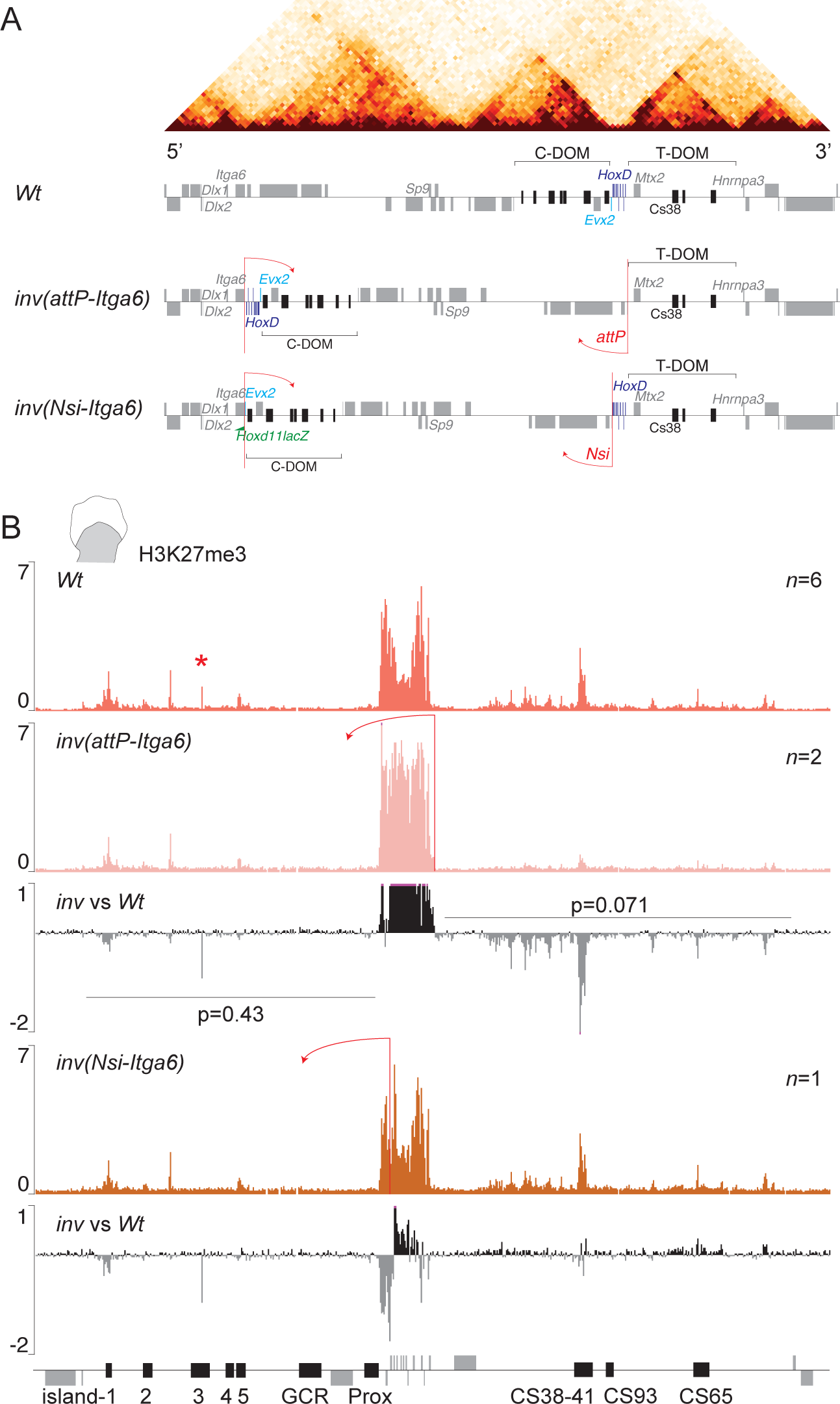
H3K27me3 distribution after disconnecting the *HoxD* from its T-DOM regulatory landscape. (*A*) Scheme of the *inv(attP-Itga6)* and *inv(Nsi-Itga6)* mutant lines. On top, a Hi-C profile from limb cells above the distribution of genes (grey) and regulatory regions (black) (chr2:71,240,000-76,320,000). The positions of both C-DOM and T-DOM are shown by brackets. In the *inv(Nsi-Itga6)* allele, an inversion is generated between the *Itga6* and the *attP* breakpoints [42] separating the *HoxD* cluster from T-DOM. In the *inv(Nsi-Itga6)* allele, the inversion occurs between the *Itga6* and the *Nsi* breakpoints [43] and hence the *HoxD* cluster remains in contact with T-DOM. In the latter inversion, a *Hoxd11lac* transgene (green flag) is inverted along. (*B*) H3K27me3 ChIP profiles from proximal limb bud cells derived from either Wild-type, *inv(attP-Itga6)* or *inv(Nsi-Itga6)* specimens. The *n* indicates the number of replicates for each track. Below each mutant dataset, a comparison of mutant *versus* control is shown. A red asterisk in the control track indicates an artifactual signal at the position of island-3.

**Additional File 4.**
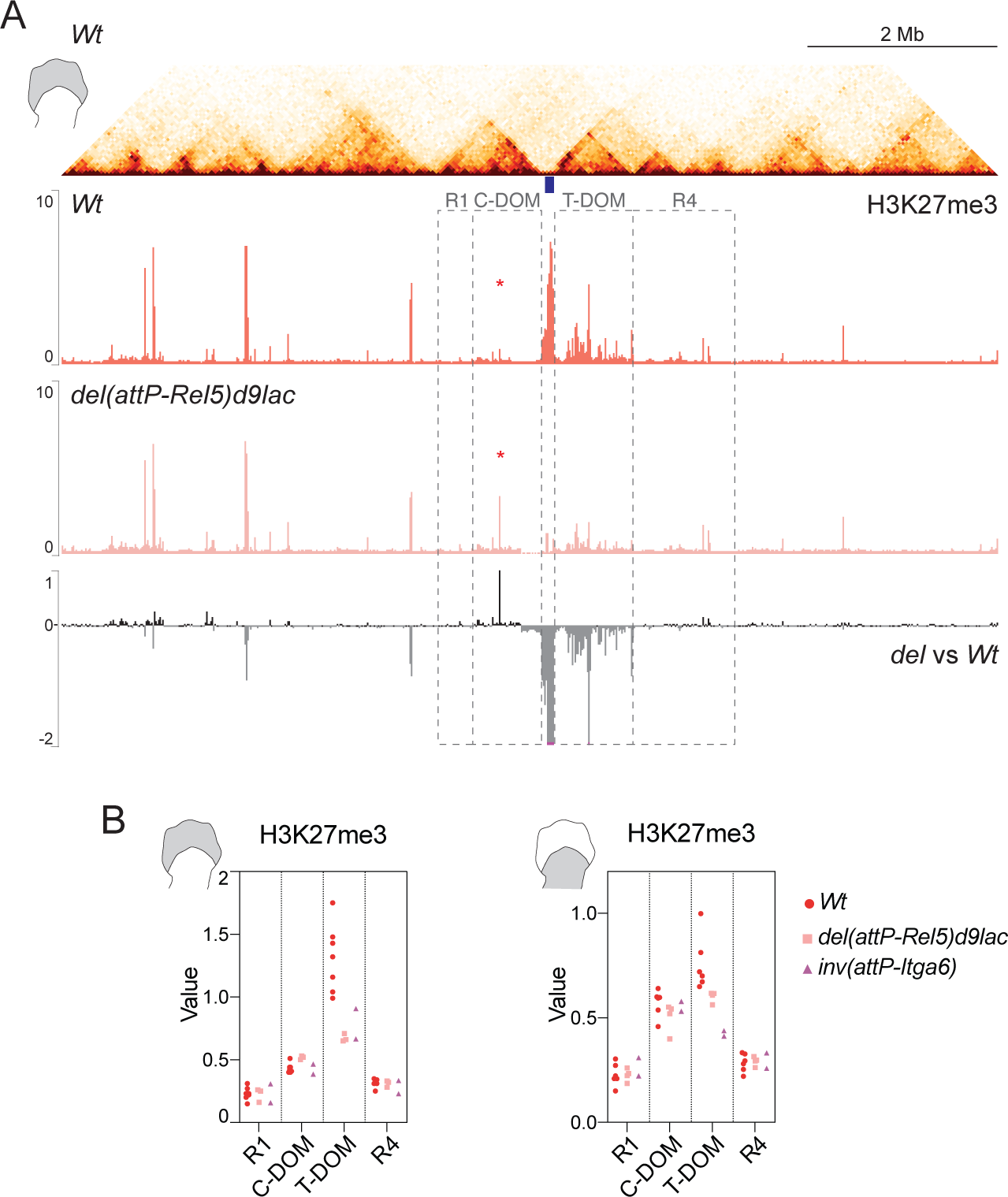
(*A*) H3K27me3 coverage outside the *HoxD* locus in distal limb bud cells from either control (*Wt*) or *del(attP-Rel5)d9lac* specimens. A corresponding Hi-C map is shown on top spanning ca. 10 megabases and centered around the *HoxD* cluster (blue box), with the related TAD structures (chr2:69,600,001-79,440,000). The flanking C-DOM and T-DOM TADs are indicated. H3K27me3 ChIP profiles in control cells show a global coverage outside the *HoxD* cluster precisely restricted to the T-DOM TAD. In *del(attP-Rel5)d9lac* mutant distal limb cells, the enrichment is much weaker. Below is the difference in the ChIP datasets comparing mutant *versus* control signals. The red asterisks point to an artifactual signal. (*B*) Quantification of H3K27me3 ChIP signal of Wild-type, *del(attP-Rel5)d9lac* and *inv(Nsi-Itga6)* in distal (left) or proximal (right) forelimb cells. The plotted values are computed from the regions depicted in (A) as dashed boxes.

**Additional File 5.**
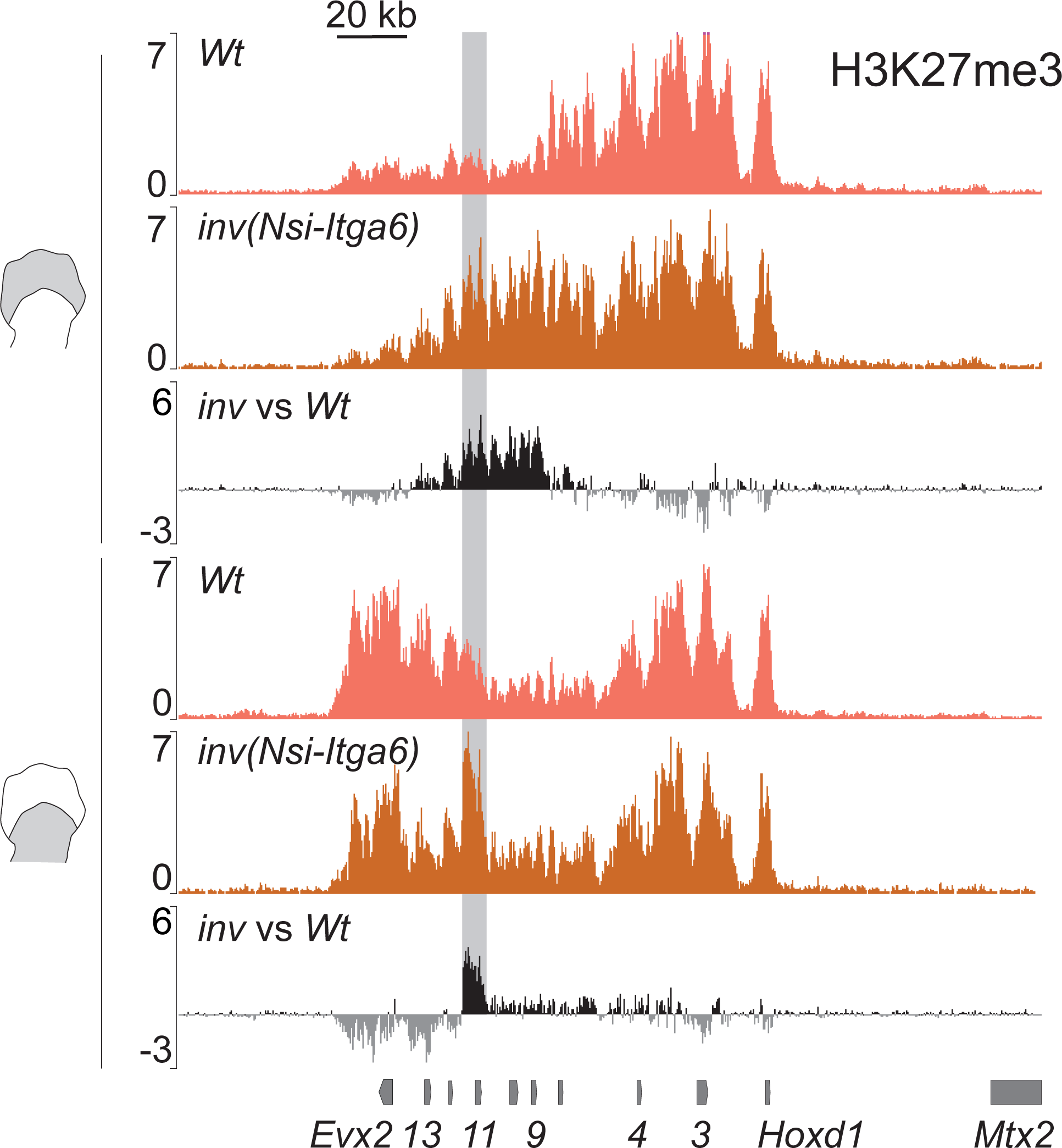
H3K27me3 signal over the *HoxD* cluster in the absence of C-DOM. H3K27me3 ChIP profiles in distal (top two tracks) and proximal (bottom two tracks) limb bud cells, either in control (*Wt*) or in *inv(Nsi-Itga6)* mutant specimen. Below are shown the difference profiles. The increase of signal in *Hoxd11* represents reads coming from the *Hoxd11lac* transgene included in the inversion allele (represented by a shaded grey box).

## METHODS

### Animal experimentation and mouse strains

Genetically modified mice were kept on a (Bl6XCBA) background and crossed in heterozygosis. Distal and proximal forelimb tissues were dissected and processed from E12.5 mouse embryos. All mutant mice used in this study and their genotyping strategies have been previously described in [34, 38, 42]. Homozygous mutant embryos were obtained by crossing heterozygous mice.

### 3D Modeling of Hi-C datasets

Hi-C original datasets from wild type, *HoxD*^*del(1-13)d9lac*^ and *HoxD*^*del(attP-Rel5)d9lac*^ were obtained from [34] (GEO accession: GSE101715). Three-dimensional modeling of the normalized 40kb binned Hi-C matrices was performed by means of the model_and_analyze.py script from the TADbit v0.2.0.58 software [36] in chr2: 73800001-75760000 (wild-type coordinates mm10). We generated 500 models for optimization and 5000 for modelling and we did not filter out matrix columns showing no interactions. We visualized the model with TADkit using the Virtual Research Environment (https://vre.multiscalegenomics.eu). Region CS38-41 (wild-type coordinates in mm10, chr2:75120051-75165771) was used as a reference mark in the 3D reconstructed Hi-C model.

### RNA extraction, RNA-seq and qPCR

Limb tissue was dissected, placed in RNAlater (Invitrogen) and directly frozen at - 80°C until further processing. After genotyping, RNA was extracted from individual samples using RNAeasy Micro Kit (QIAGEN). For RNA-seq, a total of three different biological replicates per tissue and genotype were processed. Libraries were prepared from 100ng of total RNA using the TruSeq Stranded mRNA protocol and sequenced on a HiSeq 2500 machine (100-bp reads, single end). Sequencing data were treated using the facilities of the Scientific IT and Application Support Center of EPFL. The gtf file used for STAR and cufflinks, based on Ensembl version 94 annotations, is available on figshare [58]. Adapters were removed using cutadapt (v1.16) and aligned onto mm10 using STAR [59] with ENCODE parameters. DESeq2 analysis was performed with default parameters and genes with absolute log2 fold change above 1.5 and p-value below 0.05 were considered as significant ([60] version 1.22.1). For *HoxD*^*del(attP-Rel5)d9lac*^, three biological replicates were used for each genotype and tissue. For *HoxD*^*inv(attP-Itga6)*^ only one sample was used per tissue and genotype. Track profiles show the mean of the coverage of uniquely mapped reads normalized to the number of uniquely mapped reads. They were obtained with the UCSC browser. For qPCR, purified RNA was retrotranscribed with Promega GoScript Reverse Transcriptase (Promega). Custom SYBR probes were used for quantitative real-time PCR (qPCR) in a QuantStudio 5 384-well block machine. Island3 primers were Forward: TTCCATCACAGGAGAGTCGTTG and Reverse: AGGTGGGAACATGGACTGAAAG. All other primers were described in [37, 61].

### 4C-seq experiments

The limb samples used in this study were dissected from E12.5 forelimb buds for all wild type and mutant lines. Samples were processed as in [34]. Briefly, cellular suspensions were filtered and fixed using a 2% Formaldehyde/10% FBS/PBS solution for 10 minutes. NlaIII (NEB) was used as a first cutter and DpnII (NEB) as a second cutter. DNA libraries were prepared using twelve to fourteen independent PCR reactions with 100ng DNA on each. Sequencing was performed by multiplexing several independently barcoded viewpoints. 4C-seq data were analyzed using the HTSstation web interface [62]. They were normalized to the distribution of reads on a 10Mb window and profiles were smoothened using a window of 11 fragments. 4C-seq data from wild type tissue was taken from GEO (GSE101717). Data for the CS38 viewpoint were taken from GSM2713679 and for the *Hoxd4* viewpoint from GSM2713671 and GSM2713672.

### Chromatin immuno-precipitation (ChIP)

For all samples, limb tissues were dissected and directly fixed with 1% formaldehyde in PBS for 10 minutes at room temperature, followed by 3 minutes incubation with Stop Solution from the ChIP-IT High sensitive kit (Active Motif). Samples were then washed 3 times with working Washing Solution (ChIP-IT, Active Motif) and then snap-frozen in liquid nitrogen and stored at −80°C until further processing. After genotyping, samples were pooled according to the required cell number. The total amount of tissue used for each line was different due to the size variations of the limb buds. Limb tissues were disrupted with a polytron device, lysed in RIPA buffer or Prep Buffer (ChIP-IT, Active Motif) and sonicated in Diagenode Bioruptor Pico. All H3K27ac ChIP experiments were processed as ChIP-seqs using the reagents from ChIP-IT High Sensitive kit (Active Motif). IPs were performed in parallel technical duplicates with 11 to 14µg of chromatin on each. Antibody incubation was performed overnight on a final volume of 1.5-2ml dilution buffer (0.1% SDS, 50mM Tris-HCl pH8, 10mM EDTA pH8 and proteinase inhibitors), including 2µl of H3K27ac antibody (Diagenode C15410196) at 4°C on a rotating platform. Agarose beads were added for 3 to 4h at 4°C. Washes were performed on column and DNA purification was carried out by phenol-chloroform extraction. The technical replicates were merged and yielded 1.5 to 2ng of chromatin, which were used to generate DNA libraries using the TruSeq ChIP library preparation kit. RING1B ChIP experiments were processed as for ChIP-seq using 4µl of RING1B antibody (Active Motif 39664) and following the protocol described in [32].

All H3K27me3 and EZH2 ChIP were performed following the ChIPmentation protocol [63]. Around 0.1 to 0.4 million cells were used for each IP on a final volume of 800 to 1000µl of RIPA-LS buffer (10mM Tris-HCl pH8, 140mM NaCl, 1mM EDTA pH8, 0.1% SDS, 0.1% sodium deoxycholate, 1% Triton x-100 and proteinase inhibitors), to which 2µl of H3K27me3 (Millipore 17-622) or EZH2 (Diagenode C15410039) antibodies were added. Samples were incubated for at least 2 hours with Dynabeads Protein A (Invitrogen 10001D) rotating at 4°C. Washes were performed as follows: two times RIPA-LS, two times RIPA-HS (10mM Tris-HCl pH8, 500mM NaCl, 1mM EDTA pH8, 0.1% SDS, 0.1% sodium deoxycholate, 1% Triton x-100 and proteinase inhibitors), two times RIPA-LiCl (10mM Tris-HCl pH8, 250mM LiCl, 1mM EDTA pH8, 0.5% NP-40, 0.5% sodium deoxycholate and proteinase inhibitors) and once with 10mM Tris-HCl pH8. Beads were resuspended in 24µl of tagmentation buffer (10mM Tris pH8, 5mM MgCl_2_, 10% dimethylformamide) and 1µl of Tn5 transposase (Illumina 15027865, from Nextera DNA Library Prep Kit 15028212) and transferred to PCR tubes, which were then incubated at 37°C for five minutes in a thermocycler. Samples were then resuspended and washed twice in 1ml of RIPA-LS and twice in 1ml TE buffer (10mM Tris-Hcl pH8, 1mM EDTA pH8). Beads were magnetised, DNA was eluted in ChIP elution buffer (10mM Tris-HCl pH8, 5mM EDTA pH8, 300mM NaCl, 0.4% SDS) with 2µl of proteinase K (20mg/ml stock) and then incubated for 1 hour at 55°C and 6 hours to overnight at 65°C. After de-crosslinking, the supernatant was recovered and beads were resuspended again in 19µl ChIP elution buffer with 1µl of proteinase K and left 1 hour at 55°C. The two supernatants were combined and purified with MinElute kit (Qiagen) in 22µl of EB buffer. Relative quantitation was performed using SYBR-green (as in [63]) using 2µl of DNA. Libraries were amplified according to the Cq values obtained in the previous step (12 to 14 cycles for both sets of samples), purified using Agentcourt AMPureXP beads (Beckman Coulter A63880) and eluted in 15µl of water. DNA sequencing was performed in HiSeq 2500 or HiSeq 4000 machine as 50bp single reads or 100bp single reads.

### ChIP analysis

Analyses were performed using the facilities of the Scientific IT and Application Support Center of EPFL. Sequences were filtered and adapters were removed using cutadapt (v1.16) [64] with parameters -m 15 -q 30 -a CTGTCTCTTATACACATCTCCGAGCCCACGAGAC for ChIPmentation and -a GATCGGAAGAGCACACGTCTGAACTCCAGTCAC for ChIP-seq. Reads were mapped on mm10 using bowtie2 (v2.3.4.1) using default parameters [65]. Only reads with a mapping quality above 30 were kept. A profile was obtained with macs2 [66] (version 2.1.1.20160309 option –extsize 300). Bedgraphs were normalized to their number of million tags used in the profile and replicates were merged using the tool unionbedg (bedtools v2.27) [67]. Profiles were loaded in the UCSC browser with windowing function as mean. The difference profiles were calculated using unionbedg. In order to quantify the gain or loss of chromatin marks in the C-DOM, T-DOM, and in the CS38-41 region, the number of reads falling into their respective intervals (chr2:73914154-74422050 for C-DOM, chr2:74781516-75605516 for T-DOM, and chr2:75120051-75165771 for the CS38-41 region) were assessed after duplicate removal by picard (http://broadinstitute.github.io/picard/ version 2.18.14) using the multiBamSummary function from deeptools [68]. For the C-DOM, the reads falling into the region of the artifactual peak, which is due to a PCR contamination (chr2:74207282-74208158), were excluded in all datasets. The counts were normalized to the number of reads in the input bam file and the significance was assessed by the function wilcox.test in R (https://www.R-project.org).

### Whole-mount *in situ* hybridization (WISH) and beta-galactosidase staining

Island3, *Hoxd4, Hoxd8, Hoxd9, Hoxd11, Hoxd13* and *Evx2* WISH were performed following the protocol described in [69]. The DNA fragment for Island3 probe was amplified from purified genomic DNA using primers GCAGGAATGACAGACAGGCA (Fw) and ACAGAGGTGGGAACATGGAC (Rv) and cloned into pGEM-T easy vector (Promega A1360). Beta-galactosidase staining was performed as in [34]. *Hoxd4, Hoxd8, Hoxd9, Hoxd11, Hoxd13* and *Evx2* probes were as in [70].

## ABBREVIATIONS

TAD: Topologically Associating Domain
PRC: Polycomb Repressive Complex
WISH: Whole-mount In Situ Hybridization
ChIP: Chromatin Immunoprecipitation
eRNA: enhancer RNA
lncRNA: long non-coding RNA
C-DOM: centromeric TAD
T-DOM: telomeric TAD.

## DECLARATIONS

### Ethics approval and consent to participate

All experiments were performed in agreement with the Swiss law on animal protection (LPA), under license No GE 81/14 (to DD).

### Consent for publication

Not applicable

### Availability of data and material

The datasets generated and analyzed for this study are available in the GEO repository under accession number GEOXXXX. Publicly available data used in this paper can be found in GEO under the accession numbers GSE101715 (Hi-C) and GSE101713 (4C-seq).

### Competing interests

The authors declare no competing financial interests.

### Funding

This work was supported by funds from the École Polytechnique Fédérale (EPFL, Lausanne), the University of Geneva, the Swiss National Research Fund (No. 310030B_138662) and the European Research Council grants System*Hox* (No 232790) and Regul*Hox* (No 588029) (to D.D.). Funding bodies had no role in the design of the study and collection, analysis and interpretation of data and in writing the manuscript.

### Author’s contributions

Design of experiments, ERC, DD; Bench work, ERC, NYK; Computing analysis, LLD, AUA; Analysis of results, ERC, LLD, DD; Manuscript writing, ERC, DD; Manuscript correction, LLD, NYK; Funding acquisition, DD.

#### Acknowledgements

We thank Leonardo Beccari, Aurélie Hintermann, Andréa Willemin, Jozsef Zákány and other members of the Duboule laboratories for insightful comments and discussion. We also thank Bénédicte Mascrez, Sandra Gitto and Thi Hanh Nguyen Huynh for their help with mice breeding and genotyping, as well as Mylène Docquier, Brice Petit and Christelle Barraclough from the Geneva Genomics platform (University of Geneva).

